# Vault particles are common contaminants of extracellular vesicle preparations

**DOI:** 10.1101/2023.11.09.566362

**Authors:** Xinming Liu, Zubair Nizamudeen, Christopher J Hill, Christopher Parmenter, Kenton P Arkill, Daniel W. Lambert, Stuart Hunt

**Author notes:** Correspondence: Stuart Hunt, School of Clinical Dentistry, The University of Sheffield, Sheffield, UK.

## Abstract

Extracellular vesicles (EVs) may contain a variety of molecular cargo including proteins and nucleic acids. Vault particle components have been repeatedly reported in the literature as EV cargo. Here, we demonstrated by small RNA sequencing that vault RNA (vtRNA) were highly abundant in EV pellets enriched by differential centrifugation. EVs were prepared by commonly used enrichment methods and biochemical assays used to determine whether vault particle components were bona fide EV cargo. EVs were isolated by differential centrifugation, size exclusion chromatography (SEC) and Dynabead immunocapture. RNase and proteinase treatment of EV preparations demonstrated that most vtRNA and major vault protein (MVP) were not enclosed and protected within the EV membrane. Vault-like particles were visualised in differential centrifugation pellets by cryo-transmission electron microscopy. EVs enriched by size exclusion chromatography and those isolated by immunocapture post-ultracentrifugation showed co-purification of MVP, whereas EVs isolated by direct immunocapture from conditioned medium were MVP-negative. Taken together, commonly used isolation techniques, such as differential centrifugation and SEC, can lead to contamination of EVs with vault particles. The current study highlights the importance of determining the topology of putative EV-associated components to determine if they are EV cargo or contaminants that have been co-purified.

## Introduction

Since the discovery that extracellular vesicles (EVs) can transfer functional RNA between donor and recipient cells (Valadi *et al.*, 2007; Kosaka *et al.*, 2010), there has been a massive increase in studies profiling EV RNA cargo (Batagov and Kurochkin, 2013; Crescitelli *et al.*, 2013; Huang *et al.*, 2013; Li *et al.*, 2013; Cheng *et al.*, 2014). These findings have indicated that EVs contain a large variety of RNA species including ribosomal RNA (rRNA), transfer RNA (tRNA), microRNA (miRNA), messenger RNA (mRNA), long and short non-coding RNA (ncRNA), Y RNA and vault RNA (vtRNA).

VtRNA are ∼100 nt small non-coding RNA with a stem-loop secondary structure (van Zon *et al.*, 2003). Three vtRNA paralogues are transcribed from the *VTRNA1* locus (vtRNA1-1, vtRNA1-2 and vtRNA1-3) and one from the *VTRNA2* locus (vtRNA2-1). In addition, two vtRNA pseudogenes (*VTRNA2-2P* and *VTRNA3-1P*) are annotated on the human genome assembly hg38 (Büscher, Horos and Hentze, 2020). vtRNAs account for approximately 5% of the mass of the vault particle – the largest known ribonucleoprotein complex in eukaryotic cells, localising mainly in the cytoplasm (Kedersha and Rome, 1986). Apart from the nucleic acid components, these 13 MDa subcellular organelles primarily consist of three vault proteins: major vault protein (MVP), telomerase protein component 1 (TEP1) and poly (ADP-ribose) polymerase 4 (PARP4/vPARP). MVP accounts for over 70% of the particle mass, with the outer shell of the vault containing 78 MVP copies (Tanaka and Tsukihara, 2012). Although their function has not been fully elucidated, vault particles and individual vault components have been associated with multiple cellular activities, including cytoskeleton transport, multi-drug resistance, certain signalling pathway regulation and immunity (Li *et al.*, 1999; Mossink *et al.*, 2003; Berger *et al.*, 2009; Gopinath, Wadhwa and Kumar, 2010).

Mining of the ExoCarta database revealed that MVP and vtRNA have been repeatedly reported as EV cargo or to be associated with EVs, with some studies linking them with potential intracellular trafficking and gene regulatory functions (Herlevsen *et al.*, 2007; Nolte-’t Hoen *et al.*, 2012; Lässer *et al.*, 2017). However, additional experimental work is required to assess if they are *bona fide* EV cargo. In 2014, the International Society for Extracellular Vesicles (ISEV) published a set of guidelines for EV research, which was then reviewed and updated in 2018 (Lötvall *et al.*, 2014; Théry *et al.*, 2018). MISEV2018 provided advice on using biochemical approaches to further assess the topological association of putative EV cargo to provide more convincing evidence (Théry *et al.*, 2018).

In 2019, Jeppesen *et al*. provided evidence that MVP and vtRNAs are released from cells in an exosome-independent manner, and therefore should not be considered as exosome cargo (Jeppesen *et al.*, 2019). However, other EV sub-populations were not considered. The heterogeneous nature of EVs (released from the same cell and different cell types) also raises the questions: Are vault particles a common contaminant of EV preparations? If so, how can we achieve EV isolation that is free of vaults and other similar-sized particles?

To address these questions, we compared three commonly used isolation methods (differential centrifugation, size exclusion chromatography, and Dynabead immunocapture) for enriching EVs from cancer cell line conditioned medium. We assessed the presence of EV and vault components, followed by biochemical approaches to confirm their topology/association with EVs. In the case of differential centrifugation-derived EVs, pellets collected from three increasing centrifugal speeds were assessed individually, aiming to profile the presence of vault components in pellets commonly used to enrich distinct EV subtypes. We also aimed to develop a high-purity EV isolation strategy, free from non-EV-related vault contamination. Taken together, this study explores the suitability of commonly used isolation methods in conducting EV cargo research and provides an insight into vault-free EV purification.

## Materials and methods

### Cell culture

Oral squamous cell carcinoma (OSCC) cell lines, H357 and SCC4, were routinely cultured in keratinocyte growth medium (KGM) (Allen-Hoffmann and Rheinwald, 1984) supplemented with 10% (v/v) fetal bovine serum (FBS) (Sigma). The final concentration of KGM components was: 67% (v/v) low glucose Dulbecco’s Modified Eagle’s Medium (DMEM), 23% (v/v) Nutrient Mixture F-12 Ham, 100 IU·ml^-1^ penicillin and 100 μg·ml^-1^ streptomycin, 2.5 μg·mL^-1^ Amphotericin B solution, 2 mM L-Glutamine, 1.8×10^-4^ M adenine, 0.5 μg·ml^-1^ hydrocortisone, 5 μg·ml^-1^ human insulin, and 10 ng·ml^-1^ human Epidermal Growth Factor (hEGF). Cells were maintained in a humidified cell culture incubator at 37°C with 5% CO_2_.

### Differential centrifugation

Differential centrifugation methodology was adapted from the protocol described by Théry *et al.* (Théry *et al.*, 2006). Briefly, 2 million cells were seeded in T175 flasks and allowed to adhere. After 24 h the growth medium was discarded, sub-confluent monolayers were washed in PBS, and medium was replaced with fresh KGM supplemented with 10% (v/v) ultra-filtered FBS (UF-FBS) to deplete bovine EVs as described elsewhere (Kornilov *et al.*, 2018). After 72 h, a cell confluence of approximately 80% was reached and the conditioned medium was collected for differential centrifugation. Cell debris was firstly removed by centrifugation at 300 × *g* for 10 min. The supernatant was then taken to the next centrifugation step at 2,000 × *g* for 10 min, followed by a wash with PBS and centrifugation at the same speed for another 10 min to generate a 2k pellet. Next, the supernatant was centrifuged at 10,000 × *g* for 30 min to produce a 10k pellet, followed by washing with PBS and pelleting for 30 min at the same speed. Finally, the supernatant was centrifuged at 100,000 × *g* for 1 h, to generate a 100k pellet, followed by a PBS wash and centrifugation for 1 h at the same speed. All centrifuge steps were performed at 4°C.

### Size exclusion chromatography

After 72 h, conditioned medium was collected and centrifuged at 300 × *g* for 10 min. The supernatant was concentrated to 0.5 ml using a Vivaspin-20 spin column (100 kDa MWCO) (28932363, Cytiva) by centrifuging at 6,000 × *g* for ∼45 min. Concentrated conditioned medium was fractionated by size exclusion chromatography (SEC) by application to Sepharose CL-2B resin (17014001, Cytiva) stacked in a disposable Econo-Pac column (7321010, Bio-Rad). The column was eluted with PBS and 0.5 ml fractions were collected. Fractions were stored at -20°C ready for further analysis.

### Dynabead immunocapture

Dynabeads conjugated with human anti-CD63 (10606D, Invitrogen), CD9 (10614D, Invitrogen), and CD81 antibodies (10616D, Invitrogen) were mixed at a 5:2:2 volume ratio and the mixture was used to capture EVs positive for these three tetraspanin markers, according to the manufacturer’s instructions. Mouse IgG antibody (sc-2025, Santa Cruz Biotechnology) was conjugated with M-450 Epoxy Dynabeads (14011, Invitrogen) according to the manufacturer’s protocol. Tetraspanin and mouse IgG (mIgG) beads were washed in PBS with 0.1% (m/v) bovine serum albumin (BSA) (Sigma) prior to use. For further purification of EVs isolated by differential centrifugation, EV pellets were resuspended in 1 ml PBS, combined with either tetraspanin or mIgG beads (final concentration 1.14×10^7^ beads·ml^-1^) and incubated on an orbital shaker at 4°C overnight. Alternatively, conditioned medium was centrifuged at 300 × *g* for 10 min. 5 ml of supernatant was concentrated to 1 ml using a Vivaspin-20 spin column (100 kDa MWCO) and incubated with either tetraspanin or mIgG beads. After incubation, the unbound fractions containing EVs negative for the three tetraspanins and other particles were saved and pelleted by ultracentrifugation at 100,000 × *g* for 1 h. EVs captured by the beads were firstly washed with PBS with 0.1% (m/v) BSA, then twice with PBS, followed by EV lysis for downstream analysis.

### Nanoparticle tracking analysis

Nanoparticle tracking analysis (NTA) was performed using a ZetaView PMX 110 instrument (Particle Metrix GmbH), which was calibrated with polystyrene particles of known size. 3 ml of conditioned medium or diluted EV preparations were injected into the sample cell, followed by automated acquisition and analysis by the instrument software (ZetaView 8.04.02, Particle Metrix GmbH). EV preparations were analysed using two distinct settings to detect small and large particles (Supplementary method). Particle concentrations were taken from the software generated report. Particle size and count data were taken from .txt files and used to calculate size profiles.

### ExoView analysis

EVs in conditioned medium were characterised using the ExoView R100 imaging platform (NanoView Biosciences) coupled with ExoView tetraspanin chips (NanoView Biosciences) that bind EVs positive for CD9, CD63, and CD81, with mouse IgG used as negative control. The conditioned medium was firstly centrifuged at 300 × *g* for 10 min. The supernatant was then diluted (1/2 – 1/5) in a proprietary incubation solution and loaded onto an antibody-coated chip. The chip was incubated at room temperature overnight, followed by several wash steps and the incubation with fluorescent secondary antibodies. Finally, the chip was analysed by the ExoView™ R100 reader and images were captured and analysed by the corresponding acquisition software ExoScan v0.998 (NanoView Biosciences).

### Transmission electron microscopy

EVs were imaged by the Tecnai T12 Spirit transmission electron microscope (FEI) at an accelerating voltage of 80 kV and images recorded with a Gatan orius 1000B digital camera using Gatan digital micrograph software. For resin embedded TEM imaging, EV-Dynabead complexes were fixed in fresh 2.5% glutaraldehyde in 0.1M phosphate buffer (pH7.4) overnight at 4°C, subsequently washed in 0.1M phosphate buffer twice at 15 min intervals at 4°C before being post-fixed in 2% aq. osmium tetroxide for 2 hours at room temperature, then washed again in buffer as above. Samples were dehydrated through a graded series of ethanol in water, cleared in propylene oxide and embedded in araldite resin for transmission electron microscopy. Ultrathin sections, approximately 70-90 nm thick, were cut on a Reichert Ultracut E ultramicrotome with a diamond knife and stained for 25 mins with 3% aq. uranyl acetate, washed in water, followed by staining with Reynold’s lead citrate for 5 mins. For negative staining of EVs in suspension, samples were absorbed on discharged carbon-coated copper grids for 5 min. Excess liquid was drained prior to staining with 1% phosphotungstic acid (pH 7.2) for 1 min. The grids were then washed twice in distilled water for 1 min, followed by imaging.

### Cryogenic TEM

Sample preparation for cryo-TEM was adapted from an existing protocol (Nizamudeen *et al.*, 2018, 2021). Carbon film copper TEM grids were used (EM resolutions, Sheffield, UK). Samples were left to adsorb onto the grids (5 µL/grid) for 2 min, excess solution was removed using a filter, and samples were frozen using a Gatan CP3 plunge freezing unit (Ametek, Leicester, UK), blotted for 1 s and frozen in liquid ethane. Cryo-TEM was carried out using a Tecnai Biotwin-12 at an accelerating voltage of 100 kV.

### Western blotting

EV pellets and EV-Dynabead complexes were solubilised with RIPA buffer (20-188, Millipore) supplemented with the cOmplete™ EDTA-free protease inhibitor cocktail (04693159001, Roche) and 0.1% (v/v) Pierce universal nuclease (88700, Thermo Fisher Scientific). Lysates were centrifuged at 13,000 ×*g* for 5 min and the supernatant was kept on ice for immediate use or stored at -20°C. For experiments where equal amount of total protein was loaded, protein concentrations were measured with the Pierce BCA protein assay kit coupled with the bovine serum albumin standards (23225, Thermo Fisher Scientific) according to the manufacturer’s protocol. Samples were mixed with 5× loading buffer (EC-887, National Diagnostics) and heated at 95°C for 5 min prior to the sample loading and separation by SDS-PAGE on a 10% or 12% polyacrylamide gel. Proteins were then transferred using a Trans-Blot Turbo Transfer System and the Trans-Blot Turbo Mini Nitrocellulose Transfer Packs (1704158, Bio-Rad), which were blocked with 5% (w/v) skimmed milk for 1 h at room temperature. Membranes were probed with the following antibodies (all purchased from Abcam unless otherwise stated): MVP (ab175239), TEP1 (ab64189), PARP4 (ab133745), CD9 (ab92726), CD63 (ab134045), CD81 (ab109201), TSG101 (612697, BD Biosciences), GM130 (ab52649), anti-rabbit IgG (7074, Cell Signaling Technology), anti-mouse IgG (7076, Cell Signaling Technology). For chemiluminescence detection, the SuperSignal West Pico PLUS (34580, Thermo Fisher Scientific) and WESTAR Supernova HRP detection substrate (XLS3-0100, Cyanagen) were used for high-and low-abundance proteins, respectively. Blots were scanned on a C-DiGit Blot Scanner (Li-Cor), or otherwise, exposed to a CL-XPosure film (Thermo Fisher Scientific) then developed and fixed on a Compact X4 developer (Xograph).

### Proteinase K protection assay

Differential centrifugation pellets resuspended in PBS were divided into four aliquots of equal volume and treated with: A) PBS only, B) Proteinase K (Qiagen) diluted with PBS to 20 μg·ml^-1^ final concentration, C) Triton X-100 (Sigma) diluted with PBS to 0.1% (v/v) final concentration, D) Proteinase K (20 μg·mL^-1^ final concentration) and Triton X-100 (0.1% (v/v) final concentration). All samples were then incubated at 37°C for 30 min before phenylmethanesulfonyl fluoride (PMSF, 5 mM final concentration, Sigma) was added and incubated for 10 min at room temperature to terminate proteinase digestion. Treated samples were analysed by western blotting.

### RNA extraction and quantification

RNA extraction for small RNA sequencing was carried out using the miRCURY RNA isolation kit (300110, Exiqon). RNA was extracted according to the manufacturer’s instructions. Due to unexpected discontinuity of the product, RNA extraction for other experiments was performed using the Monarch Total RNA Miniprep Kit (T2010S, New England Biolabs) according to the provided protocol.

### Small RNA sequencing

RNA sequencing was performed by the Edinburgh Clinical Research Facility (Edinburgh, UK) using the Ion Proton Platform (Thermo Fisher Scientific). Quality control checks and quantification were performed using an Agilent 2100 Electrophoresis Bioanalyzer instrument with the Agilent RNA 6000 Pico kit (Agilent Technologies), and a Qubit 2.0 fluorometer with the Qubit RNA HS Assay kit (Thermo Fisher Scientific). Using the Ion Total RNA-Seq kit v2 with an optimised protocol for low amount of short RNA cargos (<200 nt), the RNA was hybridized prior to cDNA reverse transcription and purification. cDNA was amplified with Ion Torrent adapters before the products were quantified with the Qubit 2.0 fluorometer and the dsDNA HS Assay kit while the library size distributions were obtained on an Agilent Bioanalyzer with the DNA HS kit. Equal molar quantities of libraries were combined for template preparation before sequencing on an Ion Proton instrument using a P1 v3 chip. In addition to the automatically produced BAM files (by the instrument software), microRNA reads were examined using a small RNA analysis plugin v5.0.3.0, by which the reads were aligned to mature miRNAs. Any unmapped sequences were aligned to the whole genome and counted as other RNA molecules.

### Quantitative real-time PCR

10 ng total RNA was reverse transcribed using a High Capacity cDNA Reverse Transcription Kit (4368814, Applied Biosystems) with random primers according to the manufacture’s protocol. Based on our small RNA sequencing data, three miRNAs with stable abundance across all samples (miR-23a-3p, miR-30d-5p, and miR-31-5p) were chosen to be used as internal controls. For miRNA, the TaqMan MicroRNA Reverse Transcription Kit (4366596, Applied Biosystems) was used coupled with TaqMan MicroRNA Assay 5× RT primers (miR-23a-3p 000399, miR-30d-5p 000420, and miR-31-5p 002279) to reverse transcribe 10 ng of total RNA. To quantify RNA abundance, 2× qPCRBIO Probe Blue Mix (PB20.25-01, PCR Biosystems) was combined with 20× TaqMan primers (vtRNA 1-1 Hs03676993_s1, vtRNA 1-2 Hs06632430_gH, and vtRNA 1-3 Hs04330458_s1), nuclease-free water, and cDNA (an equivalent of 250 pg RNA) for each reaction. For miRNA-qPCR, miRNA-cDNA template (an equivalent of 333 pg RNA) was used with 20× TaqMan primers specific for the above three miRNAs. qPCR was performed on a Rotor-Gene Q real-time PCR cycler (Qiagen), a two-step run was programmed: 10 min at 95°C for initial denaturation, followed by 40 cycles of 15 s at 95°C and 60 s at 60°C, in which green channel was acquired during the second step. A threshold of 0.04 was set to obtain Ct values across all experiments. Raw Ct values were then analysed using the delta-delta Ct method to generate the relative fold change of the transcript abundance compared to the selected control (Livak and Schmittgen, 2001).

### RNase A protection assay

Differential centrifugation pellets resuspended in PBS were divided into five aliquots of equal volume. Proteinase K (20 μg·ml^-1^ final concentration, Qiagen) was firstly added to the relevant samples, followed by incubation at 37°C for 30 min. 5 mM PMSF was added to all samples to halt proteinase activity by incubation at room temperature for 10 min. Triton X-100 (0.1% (v/v) final concentration) and RNase A (20 μg·ml^-1^ final concentration, Invitrogen) were added to the appropriate samples followed by incubation at 37°C for a further 30 min, before RNaseOUT Recombinant Ribonuclease Inhibitor (Thermo Fisher Scientific) was added to all samples at a final concentration of 8 U·μl^-1^ with an incubation at room temperature for 5 min. Where reagents were added to some samples, an equal volume of PBS was added to the rest. RNA was then extracted from all samples as described above.

### Data analysis and statistical analysis

Statistical analysis was performed (Prism 8, GraphPad) where data were presented as the mean ± standard deviation (SD). Data plotting was performed using Prism 8 and RStudio (RStudio). Multiple t-test with Holm-Sidak test correction were used to confirm statistical significance where applicable (p < 0.05).

## Results

### Vault particles contaminate differential centrifugation EV preparations

We firstly enriched EVs from H357 and SCC4 cell conditioned medium by differential centrifugation (DC), a method that has been used by over 80% of EV researchers (Gardiner *et al.*, 2016). The protocol was adapted from that described by Théry *et al.* in 2006 (Théry *et al.*, 2006). Conditioned medium was sequentially centrifuged at 2,000 × *g*, 10,000 × *g*, and 100,000 × *g* to generate 2k, 10k and 100k pellets, respectively.

DC-derived pellets were characterised according to the Minimal information for studies of extracellular vesicles 2018 (MISEV2018) guidelines (Théry *et al.*, 2018). Western blotting confirmed all DC-pellets were enriched with the common EV marker CD63, 2k and 100k pellets were also positive for CD9 and TSG101, respectively, whilst GM130 (an intracellular Golgi apparatus protein) was only detected in the whole cell lysate (Figure 1A). Nanoparticle tracking analysis (NTA) was used to elucidate particle concentration (Figure 1B) and size distribution (Figure 1C) of small and large particles in DC-derived pellets. Small particles were approximately 10-fold enriched in 100k pellets compared to 2k and 10k pellets. Small particles in 100k pellets had a narrower size range, with a peak diameter ∼100 nm, whilst those in 2k and 10k pellets showed broader curves that included particles of larger diameter. As expected, by analysing the same samples using an NTA setting that focused on larger particles, we were able to detect broad particle size profiles, ranging from 50 to 1000 nm. Although more large particles were detected in 100k pellets than that in 2k and 10k, these particles were mostly smaller than 500 nm. Whereas large particles in 2k and 10k pellets were enriched with those larger than 300 nm. Large particles in all pellets displayed multiple size peaks, in contrast to the single peaks observed in small particle analysis. This could indicate the heterogeneous nature of larger extracellular particles but could also result from aggregation of smaller particles. By utilising two different NTA acquisition settings we were able to gain less biased information regarding the size range of particles present in each pellet. Negatively stained transmission electron microscopy (TEM) confirmed the presence of lipid bilayer-enclosed EVs in the DC pellets, especially the 100k pellet, which contained numerous EVs with the artefactual cup-shaped morphology (Figure 1D).

**Figure 1.**
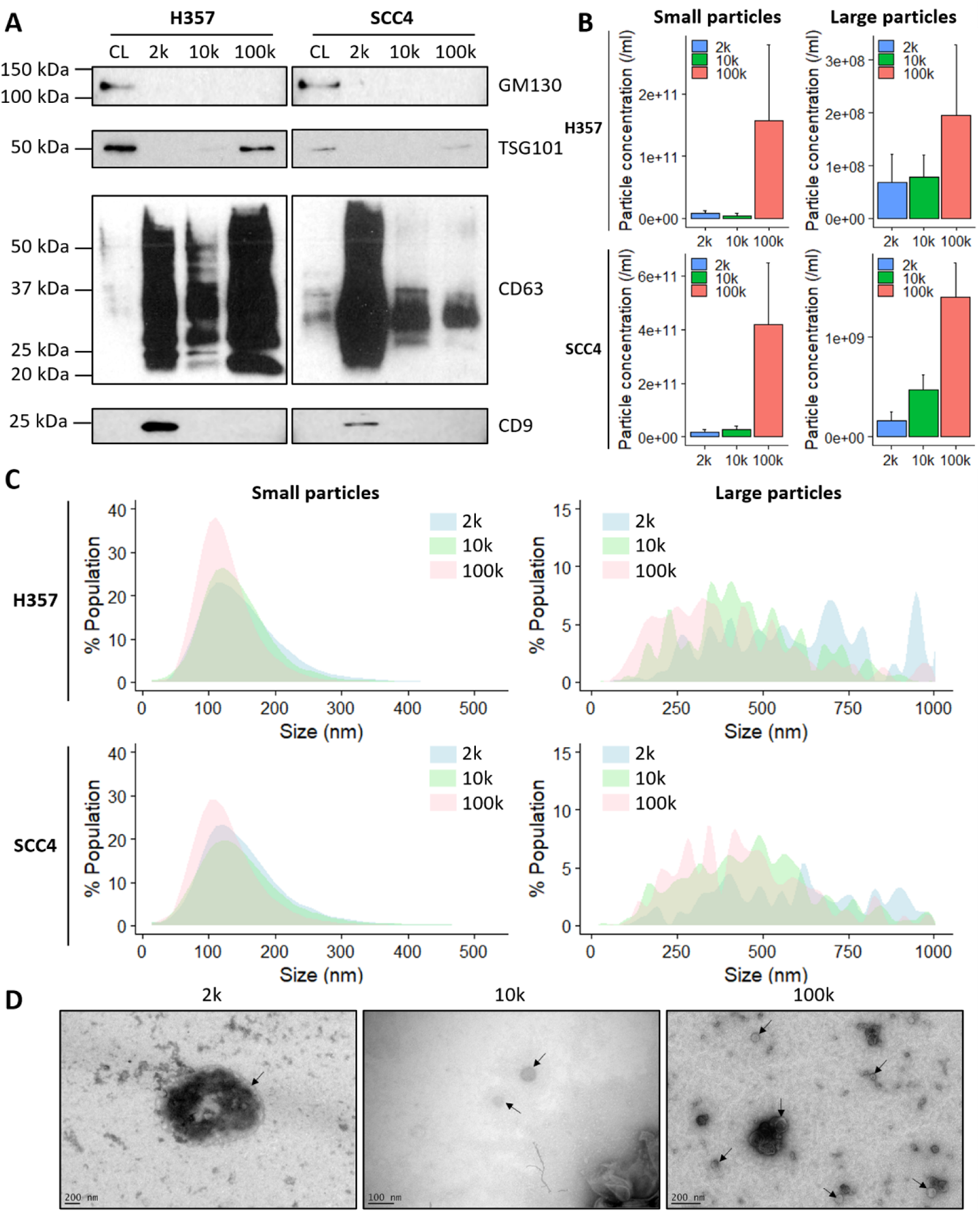
Differential centrifugation-derived pellets contain EVs. **A)** Western blots of H357 and SCC4 whole cell lysates (CL), 2,000 × *g* (2k), 10,000 × *g* (10k), and 100,000 × *g* (100k) differential centrifugation pellets. Equal quantities (2 µg) of total protein was separated by SDS-PAGE. Common EV markers (CD9, CD63, TSG101) and EV-negative marker GM130 were probed. Blots are representative of three independent repeats. **B)** ZetaView NTA showing particle numbers per ml of 2k, 10k, and 100k pellets from H357 and SCC4 cell lines. Small particles and large particles were measured using the corresponding settings on the instrument according to the manufacturer’s instructions. Data shown are mean ± SD, n=3. **C)** ZetaView NTA showing the size distribution profiles of 2k, 10k, and 100k pellets from H357 and SCC4 cells using settings focusing on small and large EVs on the instrument. Data are mean of three independent experiments. **D)** Negative stain TEM analysis of 2k, 10k and 100k pellets from SCC4 cell line. Arrows indicate EVs. Scale bars represent 200 nm, 100 nm, and 200 nm respectively.

Following confirmation that DC pellets contained EVs, we next characterised the RNA associated with 10k and 100k pellets by small RNA sequencing (Supplementary Figure 1 and Table 1), which revealed two vtRNA paralogues in the top 20 most abundant RNA species (Supplementary figure S1A). The vtRNA population was mainly dominated by vtRNA1-1, followed by vtRNA1-2 and a low abundance of vtRNA1-3, with all paralogues enriched in 100k pellets (Supplementary figure S1B). VtRNA abundance in 2k, 10k and 100k pellets was determined by qPCR, which showed a similar pattern to small RNA sequencing data (Figure 2A). We also detected major vault protein (MVP), the predominant protein component of the vault particle (Kedersha *et al.*, 1990), in DC pellet lysates by western blotting (Figure 2B).

Therefore, we assessed whether the vault components present in DC pellets were bona fide EV cargo by biochemical assays. From this point, all experiments were carried out using the SCC4 cell line due to the higher concentration of extracellular particle released (Figure 1B), hence higher protein and RNA yields for downstream experiments. We firstly utilised an RNase protection assay coupled with qPCR to determine if vtRNA present in DC pellets were protected by an EV membrane (Figure 2C and Supplementary Figure S1C). vtRNA abundance was reported relative to the average abundance of three miRNA (miR-23a-3p, miR-30d-5p and miR-31-5p) that showed abundant reads across all samples analysed by small RNA (Supplementary Table S2). The data from treatment of 100k pellets was the most straightforward to interpret, most likely due to the enrichment of vtRNA in these samples (Figure 2A). RNase A treatment alone of 100k pellets was not sufficient to degrade vtRNA1-1. However, pre-treatment of 100k pellets with proteinase K, followed by RNase resulted in a significant decrease in vtRNA1-1 abundance (Figure 2C). Treatment of 100k pellets with Triton X-100 detergent and RNase caused a significant increase in relative vtRNA1-1 abundance, which is likely due to the selective degradation of the control miRNA that were protected by an EV membrane (Supplementary figure S1D). Incubation of 100k pellets with proteinase K, Triton X-100 and RNase A resulted in insufficient RNA remaining in most samples for qPCR analysis. Similar results were also observed for vtRNA1-2 and vtRNA1-3 (Supplementary Figure S1C). Taken together these data suggest that the majority of extracellular vtRNA are protected by protein-shelled structures, rather than an EV membrane.

Similarly, we tested if the main vault structural component, MVP, was protected by an EV membrane. Proteinase treatment of 10k and 100k pellets in the absence of detergent revealed that MVP was completely digested and not protected by an EV membrane (Figure 2D). MVP in 2k pellets was not completely digested in any condition tested, which may be due to the large pellets produced (Figure 2D). In contrast, TSG101, a core component of the ESCRT-I complex, a commonly accepted intraluminal EV marker (Katzmann, Babst and Emr, 2001; Théry *et al.*, 2018), was only digested by proteinase in the presence of the membrane permeabilising detergent Triton X-100 (Figure 2D).

We next examined 100k pellets by cryo-TEM to determine if intact vault particles were present. We observed barrel-shaped vault-like particles measuring 85.2 ± 9 nm × 41.6 ± 3.7 nm (mean ± SD, n=13) (Figure 3A, 3B). We also observed numerous vesicular structures in the preparation. However, vault-like particles were not found to be physically associated with nor within EV structures (Figure 3C).

**Figure 2.**
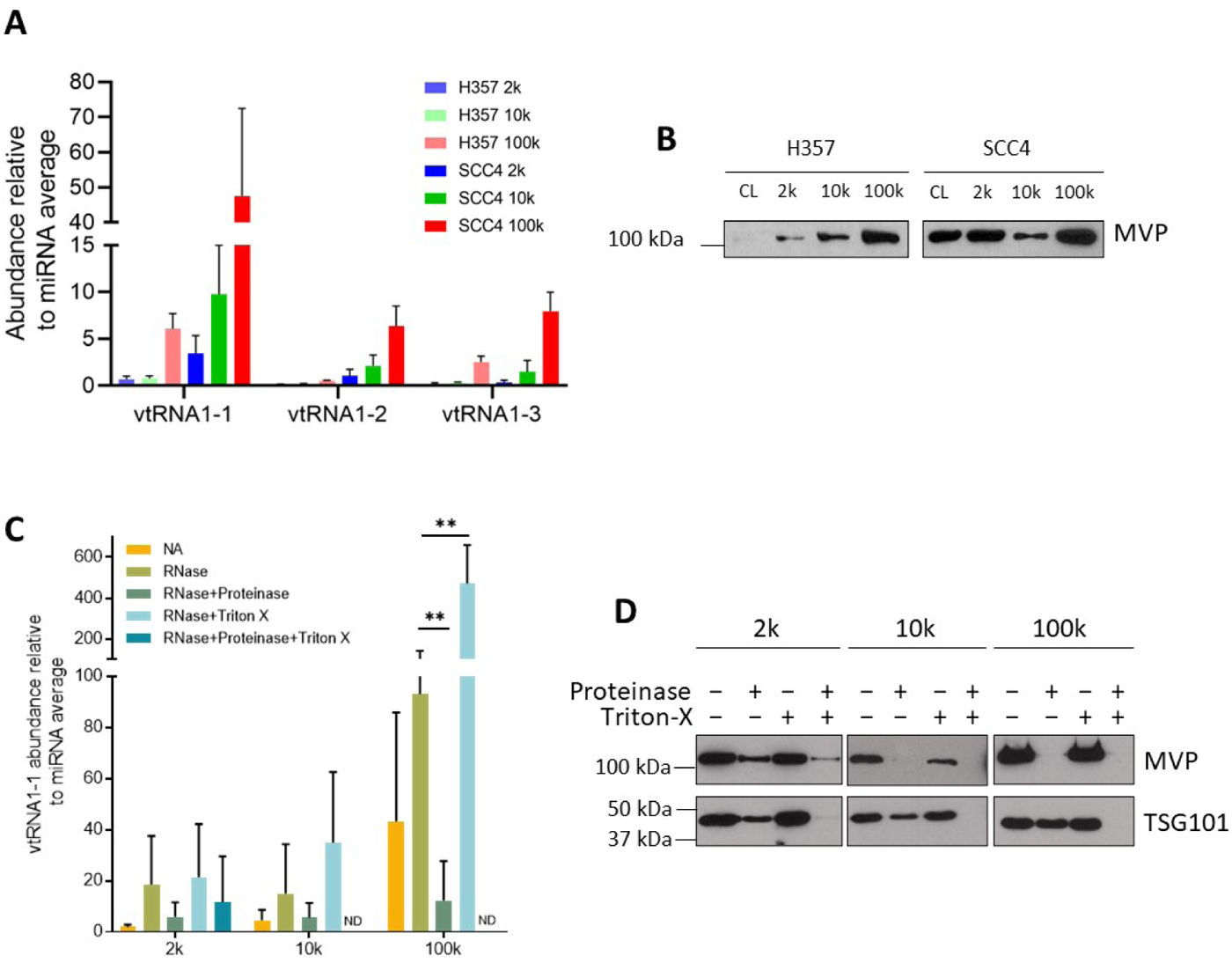
Vault components are not protected by an EV membrane. **A)** qPCR analysis of vtRNA abundance in 2k,10k and 100k DC pellets derived from H357 and SCC4 cells. VtRNA abundance is reported relative to three miRNA (miR-23a-3p, miR-30d-5p, and miR-31-5p) that were chosen as endogenous controls based on small RNA sequencing data. Data are means ± SD, n=3. **B)** Western blotting of MVP abundance in cell lysates and DC pellets derived from H357 and SCC4 cells. Equal amounts of proteins were separated by SDS-PAGE. Blots are representative of three independent experiments. **C)** RNase protection assay of SCC4 DC pellets followed by qPCR to determine vtRNA1-1 abundance. Data are means ± SD, n=3 (ND=not determined due to insufficient RNA for qPCR analysis). Statistical significance was assessed by multiple t tests corrected with the Holm-Sidak method, **p<0.01. **D)** Proteinase protection assay of SCC4 DC pellets coupled with western blotting detecting major vault protein and the luminal EV marker, TSG101, upon proteinase and membrane-permeabilising treatments. Blots are representative of three biological repeats.

**Figure 3.**
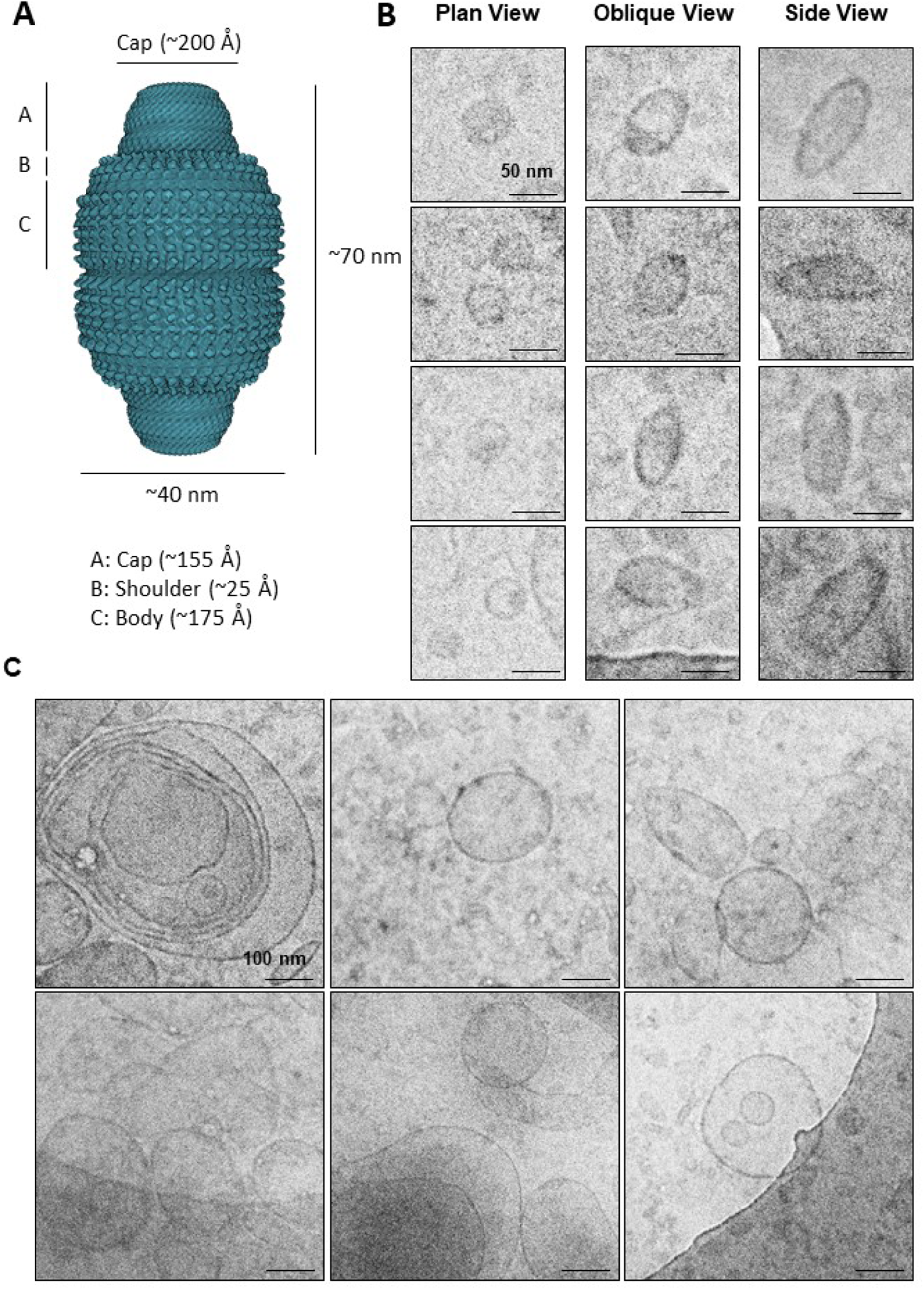
Cryo-TEM imaging of vault-like particles and EVs. **A)** Structural illustration of vault particle (PDB id: 6BP7). **B)** Collage of vault-like particles in plan (diameter = 41.2 ± 3.8 nm, mean ± SD, n=9), oblique and side view (length = 85.2 ± 9 nm, width = 41.6 ± 3.7 nm, mean ± SD, n=13). Scale bars represent 50 nm. **C)** Example images of single and multivesicular EVs ranging from 50 to 500 nm in diameter but not observed to be physically in contact with any vault-like structure (within expected size range and elliptical shape), on the EV membrane or within EV structures. Scale bars represent 100 nm.

### Vault particles co-elute with EVs by size exclusion chromatography

Having concluded that intact vault particles are present as contaminants in DC EV preparations, we assessed another well-established EV isolation technique – size exclusion chromatography (SEC). SCC4 conditioned medium was fractionated by SEC and NTA showed peak particle elution at fraction 8, with the majority of particles being enriched in fractions 7-9 (Figure 4A). Western blotting of individual SEC fractions revealed that all three vault particle-associated proteins (TEP1, PARP4 and MVP) co-eluted with EV makers (CD63, CD9 and TSG101) (Figure 4B).

**Figure 4.**
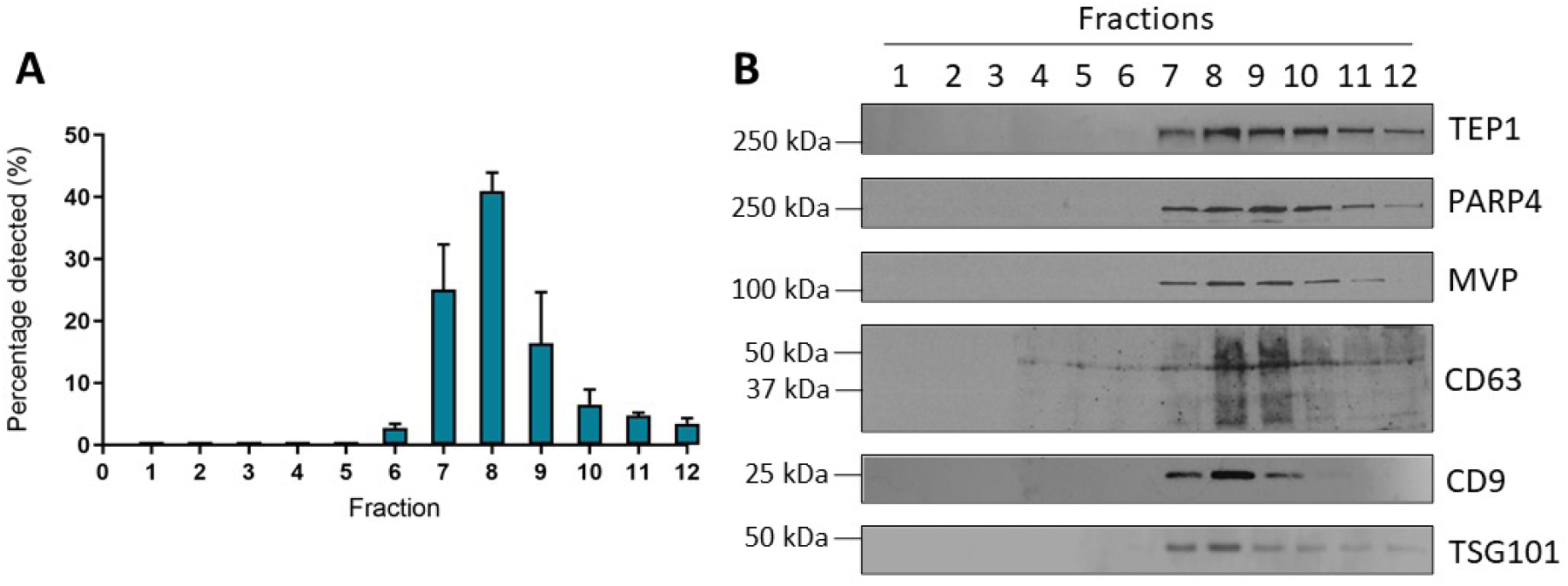
Co-elution of vault particle and EV markers by size exclusion chromatography. **A)** NTA showing SEC particle elution profile (12 x 0.5 ml fractions were collected). Bars represent mean ± SD, n=3. **B)** Western blotting to detect vault proteins (TEP1, PARP4 and MVP) and EV markers (CD63, CD9 and TSG101) in SEC fractions. Blots are representative of three independent repeats.

### A vault particle-free EV isolation strategy using immunocapture

We progressed to testing commercially available Dynabeads that capture EVs by immunoaffinity, which should reduce contamination with other similar-sized particles. The above western blotting data and ExoView analysis confirmed that SCC4 derived EVs were positive for CD9, CD63 and CD81 (Figure 5A). We therefore utilised Dynabeads to capture EVs based on this tetraspanin profile.

To determine whether immunocapture is sufficient to pull-out the marker-positive EVs from DC pellets, we applied the tetraspanin Dynabead purification to resuspended 100k pellets. EV-Dynabead complexes were examined by TEM, which showed a high level of particle aggregation at the bead surface (Figure 5B). Immunoblotting revealed that tetraspanin-positive EVs and vault particles were captured and eluted from the tetraspanin beads (Figure 5C), suggesting that particle aggregation caused by high speed centrifugation may inhibit immunoaffinity-based purification. We therefore repeated the immunocapture experiment with concentrated conditioned medium that had not been subjected to ultracentrifugation. TEM revealed the capture of individual EVs with minimal aggregation (Figure 5D). Immunoblotting indicated capture of tetraspanin and TSG101 positive EVs, but that vault particles remained unbound (Figure 5E).

**Figure 5.**
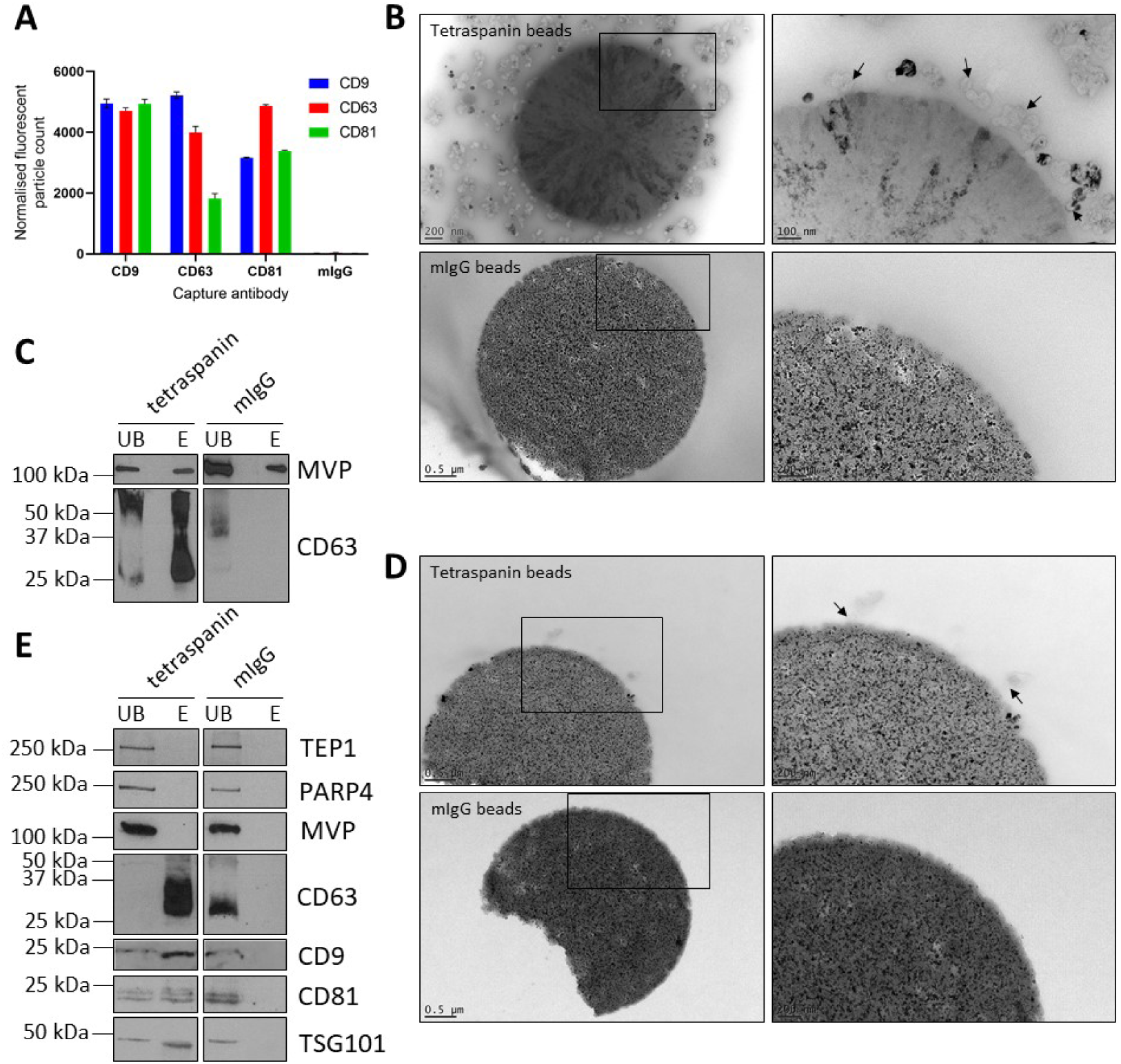
Dynabead immunocapture separates marker-positive EVs in conditioned medium from vault particles. **A)** ExoView analysis of SCC4 cell line conditioned medium using a tetraspanin microchip coated with CD9, CD63, CD81 and mIgG capture antibodies. Captured particles were labelled with fluorescent anti-CD9, CD63 and CD81 antibodies. Data are means of three technical repeats per capture antibody ± SD. **B)** Upper left: CD9/CD63/CD81 Dynabeads capturing differential centrifugation-derived 100k EV pellets with high level of aggregation. Upper right enlarged view with black arrows indicating EVs; Lower left and lower right: mIgG control Dynabeads with enlarged view. Images were obtained by negatively stained TEM. **C)** Western blot detecting MVP and CD63 in unbound and eluted fractions from tetraspanin Dynabeads and mIgG control beads after mixing with resuspended 100k EV pellets overnight. Blots are representative of three biological repeats. **D)** Upper left: CD9/CD63/CD81 Dynabeads capturing EVs from conditioned medium with no aggregation. Upper right: enlarged view with black arrows indicating EVs; Lower left and right: mIgG control Dynabeads with enlarged view. Images were obtained by negatively stained TEM. **E)** Western blot detecting vault proteins (TEP1, PARP4 and MVP) and EV markers (CD63, CD9. CD81 and TSG101) in unbound and eluted fractions from tetraspanin Dynabeads and mIgG control beads after mixing with concentrated conditioned medium overnight. Blots are representative of three biological repeats.

## Discussion

In recent decades there has been a rapid increase in EV research, with many groups attempting to determine the identity of EV-associated bioactive molecules and the effect they have upon transfer to recipient cells. Many studies utilise proteomic and transcriptomic approaches to characterise EV protein and RNA profiles, respectively. Vault particle proteins and vtRNAs have been repeatedly reported as EV-associated molecules or EV cargo (Admyre *et al.*, 2007; Buschow *et al.*, 2010; van Balkom *et al.*, 2015; Xu *et al.*, 2015). More recently, evidence has been presented suggesting that transport of MVP and vtRNAs to the extracellular space is exosome-independent (Jeppesen *et al.*, 2019).

To the best of our knowledge, the current study is the first to investigate the topology of vault particle components in EV isolates using biochemical approaches. We determined the association of vault components with EVs isolated by three commonly used techniques (Figure 6). We demonstrated that vault particles are co-isolated with EVs by DC and SEC from cell culture conditioned medium. Without further investigation, the co-purified particle components are likely to be ascribed as EV-associated molecules. Noticeably, this was identified by the MISEV2018 guidelines as one of the main issues that EV researches came across, and using biochemical approaches to further demonstrate the topological association of molecules with EVs was highly recommended (Théry *et al.*, 2018).

Vault particles measure 70 nm × 40 nm × 40 nm and hence are a similar size to small EVs. They were first discovered as major contaminants of intracellular vesicle preparations when centrifuging whole cell lysates at 100,000 ×*g* (Kedersha and Rome, 1986). In studies isolating EVs by DC and precipitation techniques, individual vault components have been repeatedly reported as EV cargo. Vault components are often detected in EV preparations enriching small EVs and exosomes (van Balkom *et al.*, 2015; Shurtleff *et al.*, 2017; Teng *et al.*, 2017). However, we also detected vault components in 2k and 10k DC pellets. This may be due to aggregation and pelleting of smaller particles subject to high centrifugal force.

**Figure 6.**
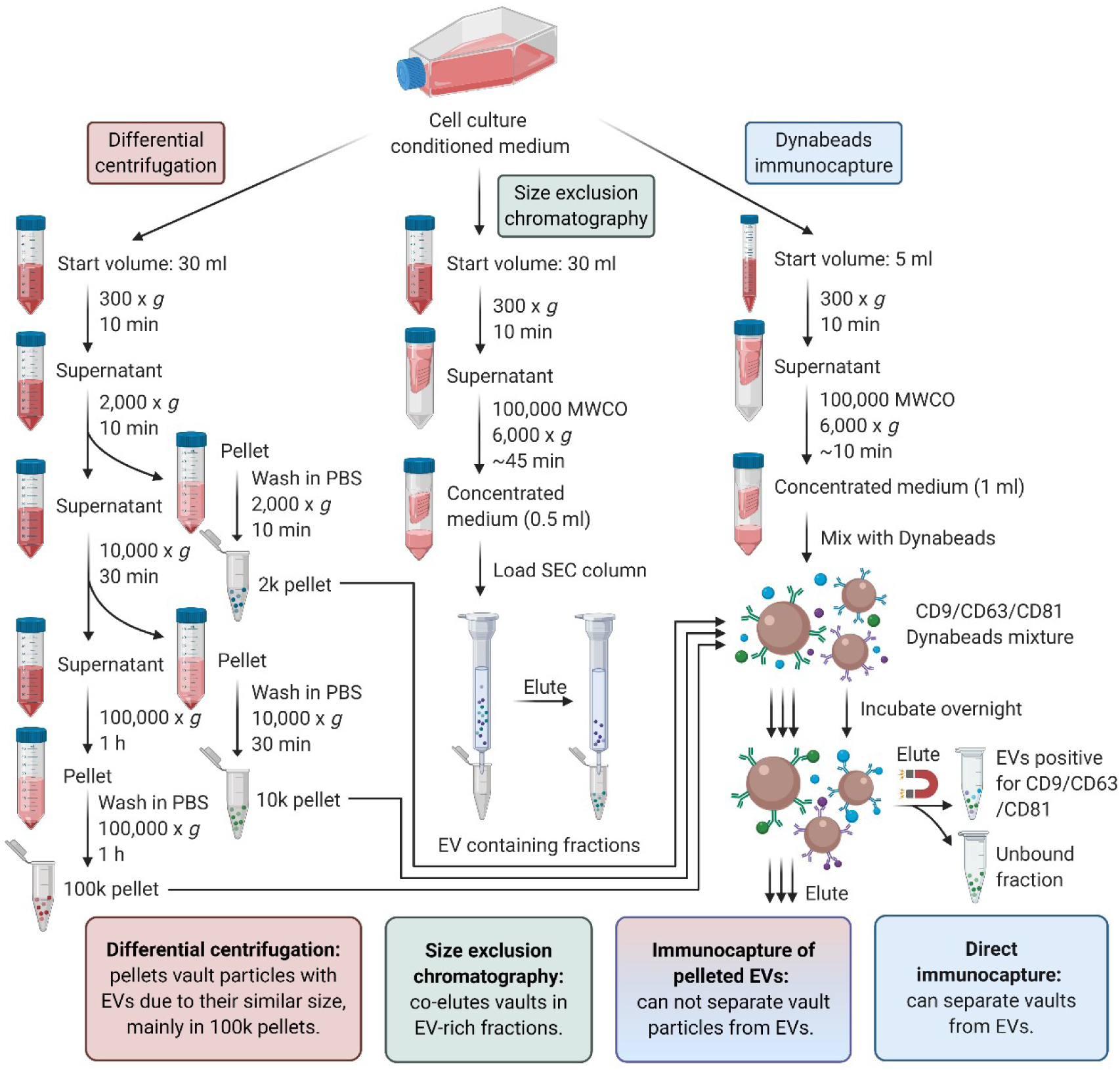
Schematic of EV isolation techniques used in this study and outcomes. EVs from cell culture conditioned medium were isolated by three techniques: differential centrifugation (DC), size exclusion chromatography (SEC), and Dynabead immunocapture. DC and SEC co-purified vault particles with EVs. Further purification of DC-derived pellets by immunocapture resulted in particle aggregation and incomplete separation of EVs from vault particles. However, EV capture from concentrated conditioned medium using Dynabeads led selection of marker specific (CD9/CD63/CD81) EVs that were free of vault particle contamination.

Proteinase protection assay data showed that MVP, the major structural component of the vault particle, was not protected by an EV membrane in 10k or 100k DC pellets. There was incomplete digestion of MVP in 2k pellets, which we speculate was due insufficient proteinase concentration or incubation time for the quantity of protein in the 2k pellets. Furthermore, the majority of vtRNAs were degraded when treating 100k pellets with RNase and proteinase (in the absence of detergent), indicating that they are mostly within a protein-shelled structure like vaults. However, there was incomplete vtRNA degradation, suggesting that some vtRNA may be protected by an EV membrane.

SEC is rapidly becoming one of the most utilised EV enrichment methods. SEC has the advantage of being relatively rapid and yielding more intact, functionally active EVs (Mol *et al.*, 2017; Monguió-Tortajada *et al.*, 2019), with some studies reporting a particle purity similar to density gradient-based isolation (Lobb *et al.*, 2015). However, SEC-derived EVs from human plasma have been shown to be contaminated with albumin and lipoproteins (Baranyai *et al.*, 2015; Stranska *et al.*, 2018). Our data suggest that vault particles are co-eluted with EVs by SEC, which is likely due to their similarity in size. Hence, researchers should be aware of the potential for SEC EV preparations to be contaminated with other non-vesicular particles.

Ultracentrifugation has been proposed as a pre-enrichment method prior to immunocapture (Pedersen, Kierulf and Neurauter, 2017). However, our data indicate that high-speed centrifuge caused particle aggregation, which prevented separation of marker-positive EVs from vaults. We were able to select tetraspanin-positive EVs using magnetic bead/antibody complexes and leave behind vaults when utilising clarified conditioned medium that had not undergone high speed centrifugation. However, immunocapture has the drawback of being marker-specific, therefore selecting a subgroup of EVs, with the loss of marker-negative populations. The use of Dynabead cocktails targeting multiple EV surface markers may be a prospective solution.

We observed vault-like particles when imaging 100k EV pellets derived from an OSCC cell line by Cryo-TEM. Taken together with the data from Jeppesen *et al.* (2019), who utilised colon cancer and glioblastoma cell lines, this suggests that their presence is likely to be universal rather than cell line-specific. Vault components have also been found in EV preparations derived from multiple body fluids and tissues, suggesting that their presence in the extracellular space is not an *in vitro* cell culture artefact (Admyre *et al.*, 2007; Gonzalez-Begne *et al.*, 2009; Skogberg *et al.*, 2013; Pienimaeki-Roemer *et al.*, 2015). It is tempting to speculate that vault export could be the result of a novel mechanism for extracellular secretion of large ribonucleoprotein particles.

In summary, we have provided compelling evidence that vault particles are a common contaminant of EV preparations generated using widely employed purification/enrichment techniques. We demonstrate that direct immunocapture allows selective enrichment of marker-positive EVs from as little as 5 ml of conditioned medium. The purified EVs are free of vault particles and can be subjected to downstream analysis. This methodology could also be used as a negative selection strategy to study marker-negative EV populations or non-EV extracellular particles like vaults.

## Acknowledgements

This research was supported by a joint PhD studentship from the China Scholarship Council and the University of Sheffield, and also generous consumables funding from The Get A-Head Charitable Trust. The authors would like to acknowledge access to the EM facilities in the Nanoscale and Microscale Research Centre (the University of Nottingham) and the Cryo-Electron Microscopy Facility (the University of Sheffield). Figure 3A and Figure 6 were created with BioRender.com.

## Supplementary data

**Table S1.** Summary of small RNA sequencing for the EV pellets from H357 and SCC4.

| Sample | Total reads | Aligned reads | Percent aligned | Mean read length | Genes detected | Isoforms detected |
| --- | --- | --- | --- | --- | --- | --- |
| H357 10k | 1,464,308 | 1,464,308 | 100.00% | 33.9 | 628 | 18,665 |
| H357 100k | 1,185,001 | 1,185,001 | 100.00% | 25.3 | 340 | 19,305 |
| SCC4 10k | 2,333,905 | 2,333,905 | 100.00% | 31.4 | 584 | 13,982 |
| SCC4 100k | 2,370,267 | 2,370,267 | 100.00% | 27.6 | 701 | 22,013 |

**Table S2.** A selection of miRNA reads in RNA-seq from the EV pellets from H357 and SCC4.

| Species | Reads per million |  |  |  |
| --- | --- | --- | --- | --- |
|  | H357 10k | H357 100k | SCC4 10k | SCC4 100k |
| MIR23A | 26493.943 | 32771.381 | 27787.256 | 28779.757 |
| MIR30D | 5885.797 | 6094.906 | 6378.303 | 9109.665 |
| MIR31 | 11186.157 | 10523.471 | 3712.921 | 4732.252 |

**Figure S1.**
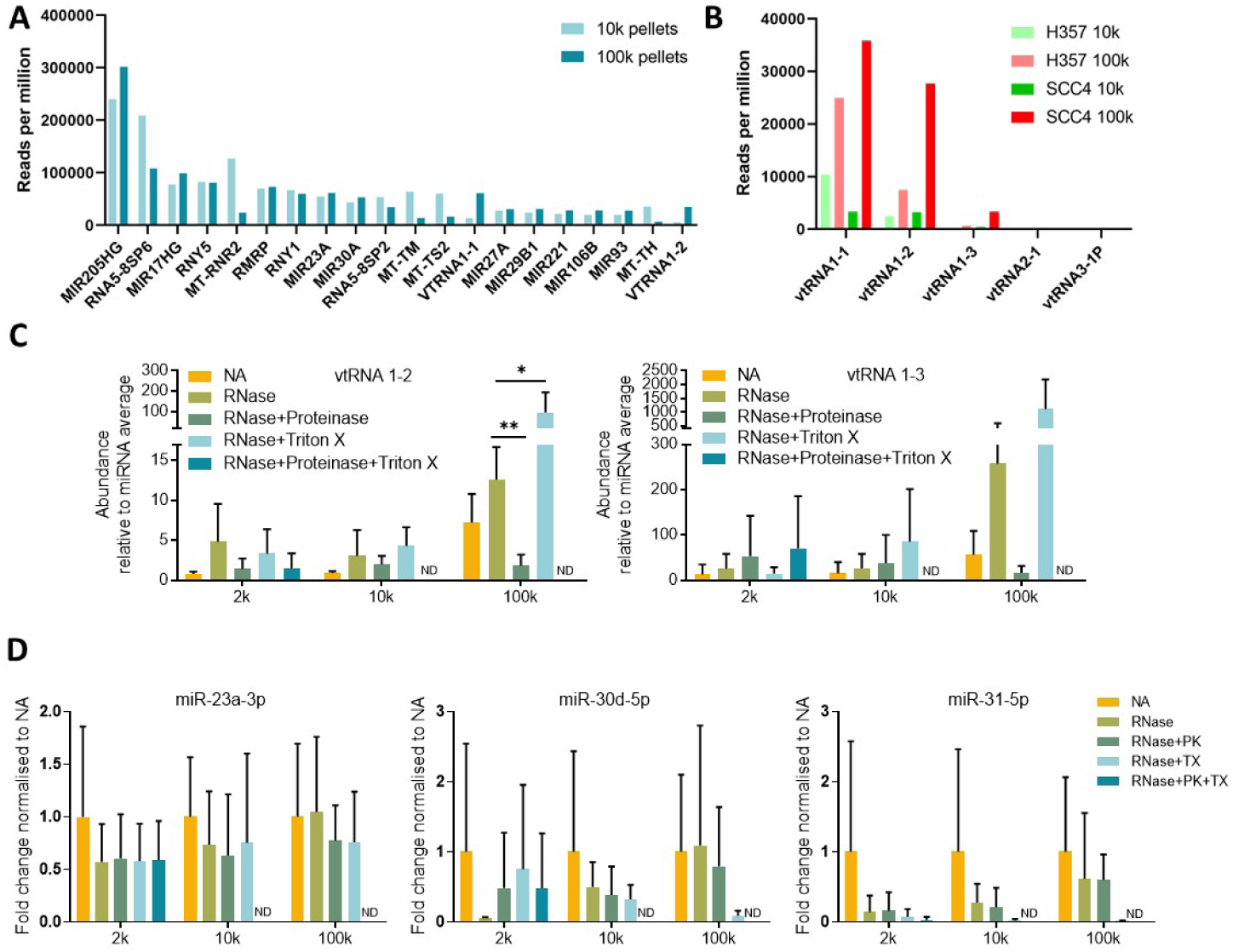
Summary of small RNA sequencing and RNase protection assay. **A)** The top 20 most enriched small RNA species in H357 and SCC4-derived DC pellets, ranked by total reads in all samples. **B)** vtRNA reads in H357 and SCC4 DC pellets determined by small RNA sequencing **C)** RNase protection assay followed by qPCR shows vtRNA1-2 and vtRNA1-3 abundance upon DC pellet treatment. Data are means ± SD, n=3, ND = not detected due to insufficient RNA material for qPCR analysis after treatment. Statistical significance was assessed by multiple t tests corrected with the Holm-Sidak method, \**p*<0.05, \*\**p*<0.001. **D)** Abundance of miR-23a, miR-30d, and miR-31 following RNase protection assay, data were normalised to the NA. Data are means ± SD, n=3, ND = not detected due to insufficient RNA material for qPCR analysis after treatment.

